# Balancing performance and complexity of dual-wedge prism-based spectroscopic single-molecule localization microscopy

**DOI:** 10.64898/2026.08.06.743389

**Authors:** Menglin Shi, Wei-Hong Yeo, Cheng Sun, Hao F. Zhang

**Author notes:** These authors contributed equally to this work (Menglin Shi, Wei-Hong Yeo).

## Abstract

Spectroscopic single-molecule localization microscopy (sSMLM) enables multiplexed super-resolution imaging by simultaneously acquiring the spatial position and spectral information of individual fluorophores. Dual-wedge prism (DWP)–based implementations provide a compact, alignment-stable approach to spectral dispersion, but trade-offs between localization precision, spectral precision, and experimental complexity remain. We systematically compare five DWP-based sSMLM configurations, including two-dimensional (2D) and three-dimensional (3D) implementations using single DWP (DWP-sSMLM) and symmetrically-dispersed DWP (SDDWP-sSMLM). We evaluate lateral precision, spectral precision, and ease of use. SDDWP configurations acquire spectral images in both channels and utilize both for spatial localization, yielding the highest lateral and spectral precision. However, for applications that do not require axial information, 2D-DWP provides a simple, plug-and-play solution with robust performance. This work offers a guideline for selecting DWP configurations based on experimental needs.

## 1 Introduction

Super-resolution microscopy techniques, including single-molecule localization microscopy (SMLM), have revolutionized our ability to visualize nanoscale structures by breaking the diffraction limit in optical microscopy^1,2^. By localizing individual fluorophores with precision beyond the resolution of conventional microscopy, SMLM enables detailed imaging of biological specimens at the molecular scale. Building on SMLM, spectroscopic SMLM (sSMLM) introduces an additional dimension by leveraging spectral information to discriminate between fluorophores, thus enhancing multiplexing capabilities and enabling dynamic molecular studies.

Reported approaches in sSMLM use diffractive optical elements, such as diffraction gratings^3–6^ or prisms^7–9^, to achieve spectral dispersion. These optical elements separate emitted light from fluorophores into spatial and spectral channels, enabling simultaneous acquisition of localization and spectral information. Despite their utility, traditional grating and prism setups often suffer from high aberrations, suboptimal transmission efficiency, and frequent optical realignment, reducing the ease of use and long-term performance.

To address these limits, we developed the dual-wedge prism (DWP) assembly to improve the performance of traditional grating and prism systems^10^. The DWP assembly exhibits lower optical aberrations, higher transmission efficiencies, and a monolithic design^11^, which simplifies its implementation into a turnkey system. By minimizing the need for continuous optical realignment, the DWP assembly provides a robust, user-friendly solution for sSMLM^12^. Recently, we designed a symmetrically dispersed dual-wedge prism (SDDWP)-based sSMLM to enhance photon utilization further and achieve superior localization and spectral precision^12^.

While these advancements have significantly improved the optical performance of sSMLM, challenges remain for fluorophores with low photon yields or those prone to photobleaching. Traditional monolithic designs introduce path length differences between spectral and spatial images, enabling three-dimensional (3D) imaging via biplane detection^10^. However, in two-dimensional (2D) imaging, such biplane configurations become unnecessary and can be counterproductive. Specifically, the path-length difference splits photons from each blinking event across two slightly defocused focal planes. As a result, the emitted signal is distributed over an enlarged point spread function (PSF) in each plane, reducing the peak intensity. This defocusing reduces the signal-to-noise ratio (SNR) and localization precision in each channel, potentially rendering weak blinking events undetectable. Consequently, the effective utilization of blinking events is reduced, ultimately limiting overall imaging quality.

To overcome this issue, we introduce the 2D dual-wedge prism (2D-DWP) configuration. By translating one right-angle prism in the DWP assembly to compensate for the path-length difference, this setup minimizes discrepancies between the spectral and spatial imaging paths, enabling photons from each blinking event to be imaged at a common focal plane. As a result, the emitted signal forms a more tightly compact PSF, increasing the peak intensity per event. This 2D configuration leads to improved SNR, enhanced localization precision, and improved detectability of weak emitters. Additionally, we explore two other relay-based optical configurations, which, although functional for 2D imaging, lack the monolithic simplicity and ease of use offered by the 2D-DWP.

In this work, we systematically compare five optical configurations in DWP-based sSMLM: 2D-DWP, 3D-DWP, 3D-SDDWP, 2D-DWP with relay, and 2D-SDDWP with relay. By evaluating their performance across localization precision, spectral precision, photon utilization, and ease of implementation, we provide insights into the trade-offs and advantages of each design, presenting a comprehensive framework for optimizing sSMLM for different needs. Here, we restrict our analysis to lateral imaging performance, as this work focuses on applications that do not require high axial precision. Accordingly, the axial precision afforded by the 3D-DWP and 3D-SDDWP configurations is not considered.

## 2. Methods

### 2.1 Imaging setup

In our imaging setup, we used a 642-nm excitation laser (2RU-VFL-P-2000-642-B1R; MPB Communications) with a filter cube (LF635/LP-C-000-NTE; Semrock), whose excitation and emission bands accommodate 642-nm excitation, together with an inverted microscope body (Ti2-E; Nikon) equipped with a 100× total-internal reflection fluorescence (TIRF) objective (CFI Apochromat TIRF 100XC Oil; Nikon). Figure 1a illustrates the emission path from the objective lens through a tube lens, with optical models of the objective and tube lens adapted from a previous study^13^. The DWP assembly is responsible for dividing the photons into a spatial imaging (0^th^ order) and a spectral imaging (1^st^ order) path, which then goes to an array detector, typically a scientific Complementary Metal–Oxide–Semiconductor (sCMOS) or an electron-multiplying charge-coupled device (EMCCD). In this work, we used a sCMOS (Prime 95B 25 mm; Photometrics) with an 11-µm pixel size for the plug-and-play setups, while we used an EMCCD (iXon Life 888; Andor) with a 13-µm pixel size for optical configurations with a relay.

**Figure 1.**
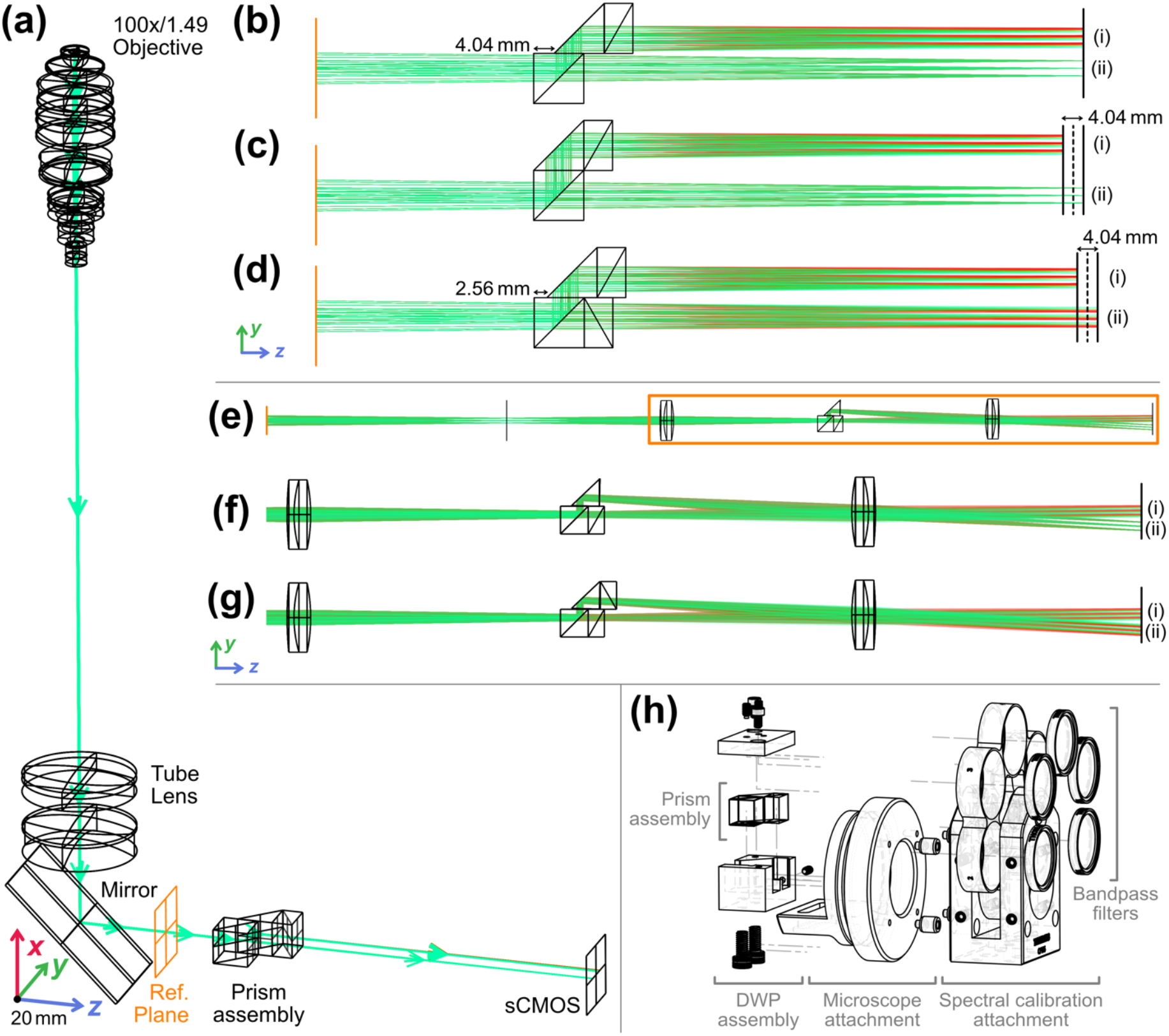
Optical configurations of 2D- and 3D-DWP-based sSMLM. (a) Emission path showing the objective lens and the tube lens to a reference plane. Plug-and-play configurations for (b) 2D-DWP, (c) 3D-DWP, and (d) 3D-SDDWP. Relay-based configurations for (e) Relay optics setup, with magnified view of (f) 2D-DWP with relay, and (g) 2D-SDDWP with relay. In configurations (b-g), (i) denotes the +1^st^-order spectral image and (ii) denotes the spatial image, except in the SDDWP configurations (d) and (g), where (ii) corresponds to the −1^st^-order spectral images. (h) Assembly drawing of mechanical parts attached to a side port of Nikon microscopes.

For plug-and-play setups, we have 2D-DWP (Fig. 1b), 3D-DWP (Fig. 1c), and 3D-SDDWP (Fig. 1d) configurations. These configurations consist of 3 key optical components: a non-polarizing beamsplitter (BS), a right-angle prism (RAP), and a DWP comprising two 30º wedge prisms made of N-LAF21 and N-SF11 glass materials bonded together to form a single dispersive element (Hyperion Optics). The BS (BS010; Thorlabs) is used to split the image into a (i) reflected and (ii) transmitted path, and in the reflected path, the RAP (84-514, Edmund Optics) is used to reflect the beam, so the resulting beam is parallel to the transmitted path. To induce spectral dispersion, a DWP is typically added to the reflected path after the RAP. We note that the non-polarizing beamsplitter and right-angle prism shown in can be combined into a lateral displacement beamsplitter, further simplifying the optical setup (which is not shown here).

The 3 key components, when placed in the 3D-DWP setup (Fig. 1e), yield a path-length difference of 4.04 mm, enabling 3D imaging via biplane detection, as previously reported^5,10,14^. By shifting the RAP and DWP in the 2D-DWP forward by 4.04 mm, we eliminate the path length difference in 3D-DWP (Fig. 1c). In the 3D-SDDWP setup, adding one DWP to the transmitted path increases the focusing position of the imaging plane of the transmitted path, but this may be compensated by shifting the RAP and DWP in the reflected path by 2.56 mm, which gives us the same path length difference at the imaging plane of 4.04 mm.

To compensate for path length variations inherent in 3D configurations, we used a pair of 100 mm lenses (49-390, Edmund Optics) which gives us two relay setups (Fig. 1e): 2D-DWP with relay (Fig. 1f), and 2D-SDDWP with relay (Fig. 1g). To ensure that the beams from the two imaging paths do not focus back to the same spot after splitting with the BS, we introduced a 1.5º tilt on the RAP, which gives us two distinct images on the imaging plane. To minimize aberrations introduced by the 100-mm relay lens, we equalized the path lengths between the reflected and transmitted paths and directed the reflected beam through the lens’s center.

Finally, for non-relay setups (Figs. 1b-1d), we designed a microscope adapter attachment (Fig. 1h), which holds the DWP assembly. The adapter provides an easy-to-use, plug-and-play setup for Nikon microscopes, though similar designs can also be adapted for other microscopes. The adapter also provides connections to the 30-mm cage mounting system, which can be used with a Thorlabs filter wheel as an additional attachment for spectral calibration^14^.

### 2.2 Spectral Calibration

The images detected by the camera are illustrated in Fig. 2a. These images appear in different combinations depending on the optical configurations summarized in Fig. 1. Specifically, the single-DWP configurations shown in Figs. 1b, 1c, and 1f produce a pair of images consisting of an 0^th^-order spatial image and a +1^st^-order spectrally dispersed image on the camera. In contrast, the dual-DWP (SDDWP) configurations shown multiplying a factorhown in Figs. 1d and 1g generate two spectrally dispersed images corresponding to the -1^st^ and +1^st^ diffraction orders.

**Figure 2.**
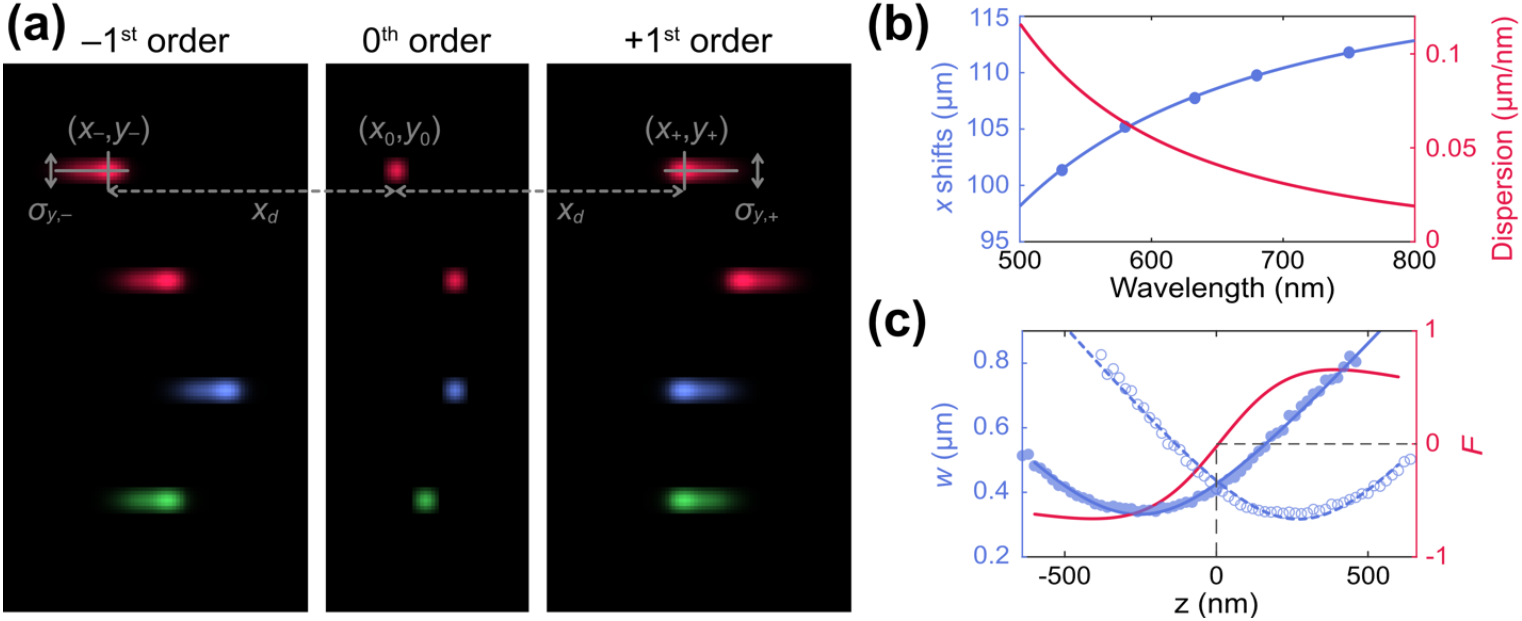
(a) Illustration of representative single-emitter point spread functions (PSFs) recorded in -1^st^, 0^th^, and +1^st^ diffraction orders. The 0^th^ order provides the undispersed spatial image at (***x***_**0**_, ***y***_**0**_), while the ±1^st^ orders yield spectrally dispersed images at (***x***_±_, ***y***_±_) with a wavelength-dependent lateral shift ***x***_***d***_ and focus-dependent PSF widths **σ**_***y***,±_. (b) Spectral calibration showing the lateral shift of the dispersed order (blue) and the corresponding dispersion (red) as a function of emission wavelength. (c) Axial calibration curve of the biplane detection for 3D imaging.

In all cases, spectral information is encoded as a wavelength-dependent lateral displacement between paired images. As illustrated in Fig. 2a, emission from a single fluorophore is recorded either as a 0^th^-order spatial image at (*x*_0_, *y*_0_) and a dispersed ±1^st^-order spectral image at (*x*_±_, *y*_±_) in single-DWP configuration; or as two symmetrically dispersed ±1^st^-order images in dual-DWP configurations. The relative lateral shift quantifies the spectral coordinate *x*_*d*_ = *x*_±_ − *x*_0_ for single-DWP setups, while in the SDDWP setup, the coordinate is quantified by the separation between the -1^st^ and +1^st^ orders where *x*_*d*_ = *x*_+_ − *x*_−_.

To calibrate the localization positions after spectral dispersion relative to known wavelengths, we used a nanohole array imaged through five bandpass filters in the spectral calibration attachment (Fig. 1h) with a fluorescent LED illumination system (TI2-D-LHLED; Nikon). The calibration protocol followed is detailed in our previous work^12^. We used bandpass filters centered at 532 nm, 580 nm, 633 nm, 680 nm, and 750 nm (FL532-3, FBH580-10, FLH633-5, FBH680-10, and

FBH750-10; Thorlabs).

### 2.3 Axial Calibration

To calibrate axial positions, we prepared fluorescent microsphere samples (F8807; Thermo Fisher) diluted approximately 10^6^ times in phosphate-buffered saline (PBS; J61196; Thermo Scientific). The resulting single-microsphere samples were imaged using a 642-nm excitation laser. Images were captured over a range of −2.0 µm to +2.0 µm relative to the focused spot in 20-nm increments. We fitted these images with two parabolas to estimate the defocus curve as shown in Fig. 2c^14^.

### 2.4 Image processing with ThunderSTORM

To process the captured images and extract meaningful localization data, we used ThunderSTORM to perform localization on these images using Maximum Likelihood Estimation (MLE) Gaussian fitting. For the spatial images, an intensity threshold of 1.2 times the standard deviation of the first wavelet level was applied for approximate localization. The localization was followed by sub-pixel refinement with a fitting radius of 3 pixels and an initial sigma guess of 1.5 pixels.

For spectral image processing, we used an elliptical Gaussian point spread function (PSF) model in ThunderSTORM. The intensity threshold was set to 1.1 times the standard deviation of the first wavelet level, with a fitting radius of 4 pixels and an initial sigma of 2.5 pixels. These parameters were empirically chosen to account for the broader PSFs observed in spectral images compared to spatial images. After completing the localization process in ThunderSTORM, we exported the coordinates and associated statistics for all localizations as CSV files for further analysis.

## 3. Results and Discussion

### 3.1 Factors affecting SMLM performance

SMLM leverages diffraction-limited optics to generate PSFs from emitters, which are captured and typically fitted with a Gaussian function to determine the emitter’s sub-diffraction positions. This process enables super-resolution performance. The Gaussian approximation of a PSF can be described by

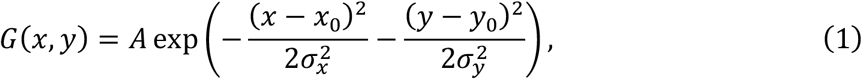

where *x*_0_ and *y*_0_ are the true *x*- and *y*-positions of the localization, and σ_*x*_ and σ_*y*_ are the standard deviations of the PSF in the *x*- and *y*-directions, respectively. The FWHM of this Gaussian fit can be obtained from σ_*x*_ and σ_*y*_ by multiplying a factor of 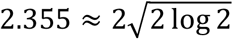. As detectors have finite pixel sizes, the intensities captured by a detector with pixel size *a* can be described as

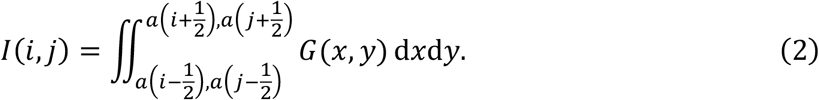

The sum of these intensities over a region of interest sufficiently large to cover most of the signal gives the number of emitted photons, *N*, which is typically limited in SMLM, as fluorophores usually have a limited photon budget before photobleaching^2^. When fitting the discretized PSF with a Gaussian function using maximum likelihood estimation, the lateral localization precision, Δ*x*, of a single localization is determined by^15^

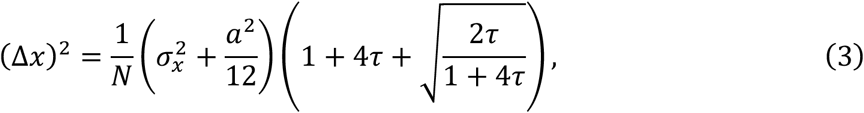

where the background correction factor, 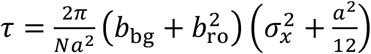 , where *b*_*bg*_ and *b*_ro_ are the mean number of background photons per pixel and the standard deviation of readout noise, respectively. From Fig. 3a, the contour map provides a comprehensive view of how localization precision depends jointly on photon count *N* and PSF width (FWHM). Along the *N*-axis, localization uncertainty decreases rapidly at low photon counts and then gradually approaches a slower rate of improvement, reflecting the 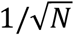 dependence in Eq. (3). Along the FWHM axis, increasing PSF width results in reduced localization precision due to the broader spatial distribution of photons. The iso-precision contour lines highlight the trade-off between these two factors: increasing photon count can partially offset PSF broadening, but only within a limited range. At large FWHM, even substantial increases in photon count yield diminishing improvements in precision.

**Figure 3.**
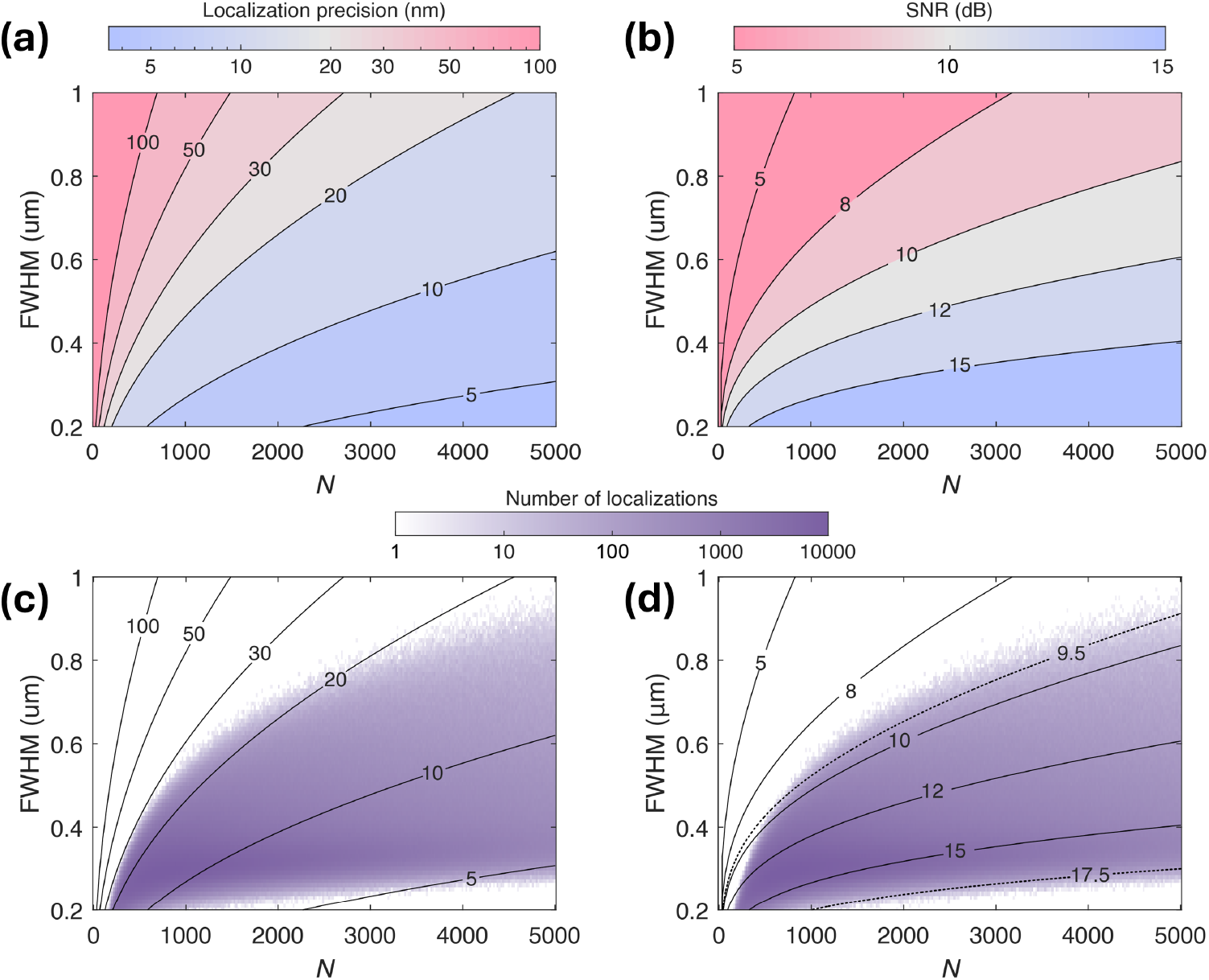
Contour maps of localization performance as a function of photon budget and the FWHM of PSF. (a) Lateral localization precision is shown as a color map over the two-dimensional parameter space of detected photon number ***N*** and PSF FWHM. Black contour lines represent iso-precision curves, indicating combinations of ***N*** and FWHM that give the same lateral precision. (b) SNR displayed as a color map over the two-dimensional parameter space of detected photon number ***N*** and PSF FWHM, with black contour lines denoting constant-SNR levels. (c) Overlay of the number of localizations from a sample image of tubulin labeled with DY-634 on the parameter space of detected photon number ***N*** vs. FWHM of PSF. The contour shows the combination ***N*** and PSF that gives the same lateral precision, which is the same as (a). (d) Overlay of the number of localizations from a sample image of tubulin labeled with DY-634 on the parameter space of the detected photon number ***N*** vs. FWHM of PSF. The contour shows the combination ***N*** and PSF that gives the same SNR, which is the same as (b).

An alternative description of SMLM performance can also be defined with an SNR as the peak intensity divided by total noise, or

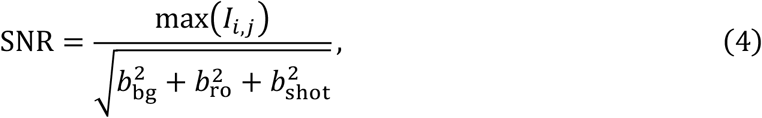

where the Poisson-distributed shot noise, 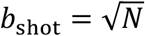, with 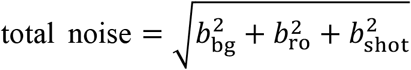 .

Assuming an infinitesimally small PSF which is less than the pixel size, we have the following simplification:

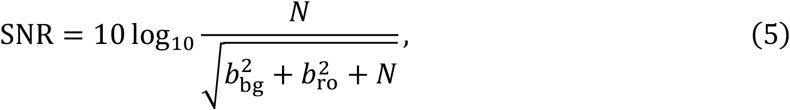

where for large *N*, the shot noise becomes the dominant noise source, and 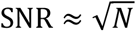 . This is evident in Fig. 3b, where the SNR isocurves follow approximately square-root curves. Increasing the width of the PSF results in a lower SNR, as the peak is less prominent with an increase in the spread of the PSF.

To gain insight into the typical localization precision and SNR achieved in practical SMLM experiments, Figs. 3c and 3d overlay the distributions of localizations from a representative image of tubulin labeled with DY-634 onto the contour maps in the two-dimensional parameter space defined by detected photon number *N* and PSF FWHM. In Fig. 3c, the isocurve represents combinations of *N* and FWHM that yield the same lateral localization precision, identical to those shown in Fig. 3a. The majority of localizations fall within isocurve corresponding to lateral precisions of approximately 5-20 nm, indicating the typical performance range for this dataset. Similarly, Fig. 3d overlays the same localization distribution onto the SNR isocurve map shown in Fig. 3b, where contour lines denote combinations of *N* and FWHM that produce the same SNR. Most localizations are concentrated in regions corresponding to SNR values between approximately 9.5 dB and 17.5 dB.

Overall, optimal SMLM performance is achieved when the detected photon count is maximized, PSF broadening and aberrations are minimized, and background noise is low. These considerations are particularly important for sSMLM with DWP, where photons must simultaneously support accurate spatial localization and spectral identification.

### 3.2 Lateral precisions of different configurations

In standard SMLM, the lateral positions of emitters are estimated by fitting a 2D Gaussian function to the PSF using maximum likelihood estimators or through least-squares fitting. With the 2D-DWP, 3D-DWP, and 2D-DWP with relay setups, this is done similarly on the spatial image (labeled (ii) in Figs. 1b, 1c, and 1f), as the imaging path is not spectrally dispersed for spectroscopy. With the SDDWP setups, as both the reflected and transmitted paths give spectral images (labeled (i) and (ii) in Figs. 1d and 1g), we fit both PSFs in each image. Thereafter, the lateral position and, subsequently, the lateral precision can be determined from the mean localization.

In 3D configurations where axial positions are provided through a path length difference, the detector is positioned in the middle (dotted line) of imaging planes (i) and (ii) in Figs. 1d and 1e. With biplane setups, the widths of the PSFs approximate two parabolas (Fig. 4a) when the emitter is defocused in the *z*-direction. Using an axial calibration procedure, the axial position can be determined, but this is beyond the scope of this work. For strictly 2D applications, we may ignore the axial positions provided and use only the lateral positions from 3D-DWP and 3D-SDDWP, as in a 2D imaging setup.

**Figure 4.**
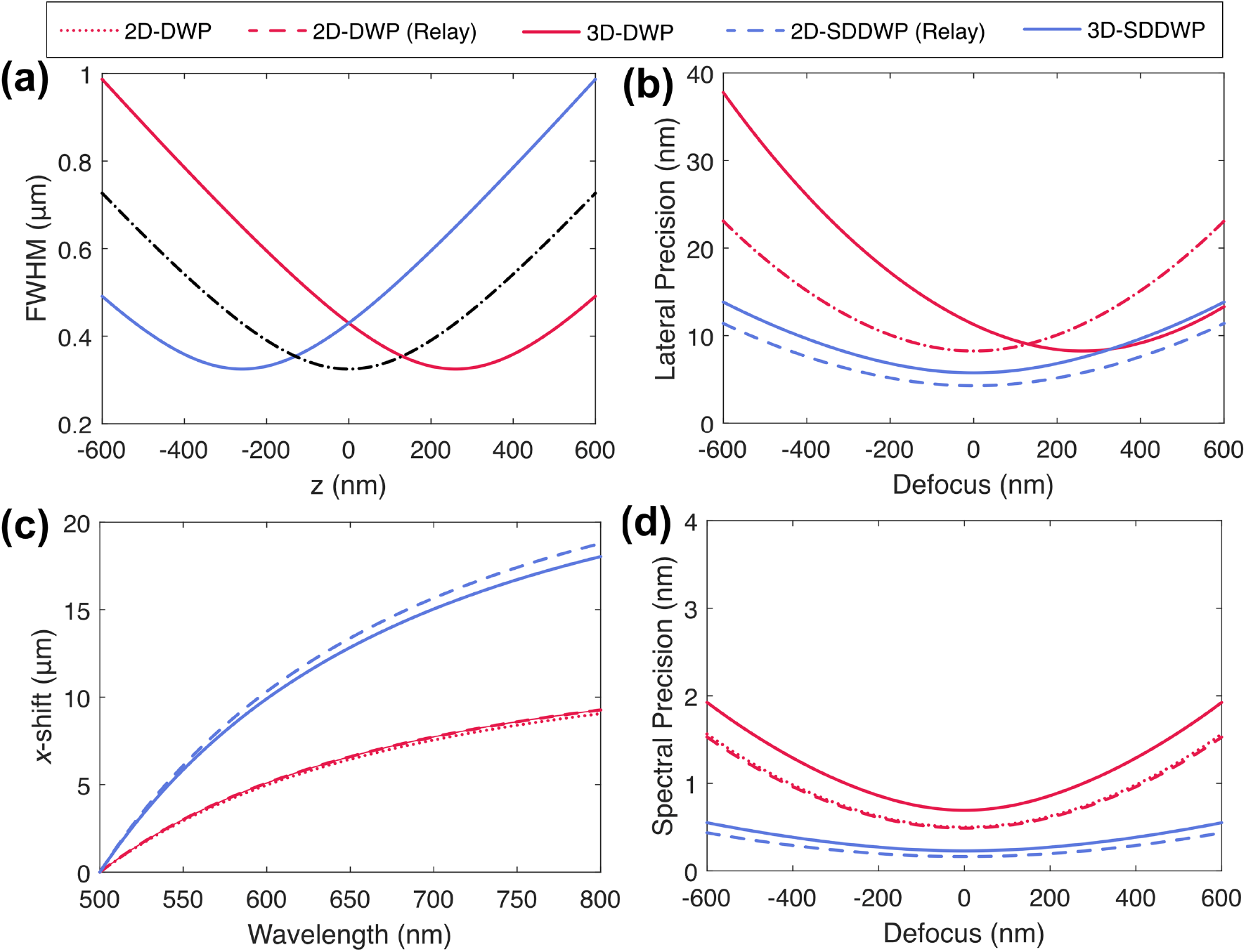
Simulated comparison of optical performance for DWP- and SDDWP-based sSMLM configurations. (a) FWHM of the PSF as a function of defocus **z** for the two imaging planes in biplane configurations, showing defocus-dependent PSF broadening. (b) Spectral calibration curves for DWP (red) and SDDWP (blue) configurations, showing increased effective dispersion for symmetrically dispersed designs. (c) Lateral localization precision as a function of defocus for 2D and 3D configurations. (d) Spectral precision as a function of defocus for all configurations, highlighting the performance advantage of SDDWP over the DWP setup.

In all configurations, the spectral identity of an emitter is determined by comparing the *y*-shifted positions between images (i) and (ii), which will be discussed in more detail in Sec. 3.3. As all configurations rely on both images (i) and (ii) during processing, having high SNR in both images maximizes the probability of detection. This point occurs with defocus = 0 nm, and from Fig. 4b, we see that SDDWP variants generally provide the best lateral precision, as they utilize photon budgets from both images (i) and (ii) for estimation of lateral positions. 3D-DWP has optimal localization precision with a defocus of 250 nm, as the smallest FWHM is obtained at that point (Fig. 4a), where the spatial image (ii) is in focus on the imaging plane. 2D-DWP and 2D-DWP with a relay are expected to perform similarly, since adding the relay should not significantly affect optical performance. Overall, when defocus is minimal, the 3D configurations generally perform worse than the 2D configurations, as defocus in the biplane setup increases FWHMs.

### 3.4 Spectral precisions of different configurations

As discussed above, the spectral identity of an emitter can be determined through the comparison of the *y*-shifted positions between images (i) and (ii), and absolute wavelengths related to the *y*-shifted positions are obtainable through spectral calibration. Although DWPs have non-linear spectral dispersion, a rational invertible empirical function, *f*(*λ*), can be used to correct its non-linearity and provide a simple linear relation between *y*-shifts, *x*_*λ*_, and *f*(*λ*) as

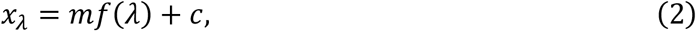

where *f*(*λ*) is the function that maps the wavelength to a corresponding scaled x-position of the nanohole array.

Fig. 4c shows the spectral calibration curves of the 5 configurations. The SDDWP configurations exhibit a larger effective separation between paired spectral images than the DWP configurations, corresponding to a higher effective dispersion over the same wavelength range. This increased separation directly improves sensitivity to wavelength differences and forms the basis for enhanced spectral precision in SDDWP-based systems.

The resulting spectral precision as a function of defocus is shown in Fig. 4d. In general, 2D configurations exhibit better spectral precision than their 3D counterparts. This is because 2D implementations operate at a single focal plane, avoiding the intentional defocus introduced in biplane (3D) imaging. In 3D configurations, defocusing increases the PSF size, which degrades the localization precision of the *x*-positions and directly propagates to reduced spectral precision. At the best focus (*z* = 0 nm), the 3D DWP configuration achieves a minimum spectral precision of approximately 0.8 nm, while the 3D SDDWP configuration reaches approximately 0.4 nm. In comparison, the corresponding 2D configurations consistently achieve lower spectral uncertainty across the entire defocus range, as shown in Fig. 4d.

Key results from simulations for the different optical configurations shown in Fig. 4 are summarized in Table 1. Among the 2D implementations, 2D-SDDWP (relay) achieves the best overall performance, with lateral and spectral precisions of 4.3 nm and 0.16 nm, respectively, reflecting the benefits of symmetric dispersion and dual-channel spatial estimation. In comparison, 2D-DWP and 2D-DWP (relay) exhibit similar lateral (∼8.3 nm) and spectral (∼0.5 nm) precision, indicating that the addition of relay optics does not significantly alter performance. For 3D configurations, performance degrades due to intentional defocus in biplane imaging. However, 3D-SDDWP substantially improves both lateral (5.8 nm) and spectral (0.23 nm) precision relative to 3D-DWP (11.3 nm and 0.69 nm).

**Table 1.**
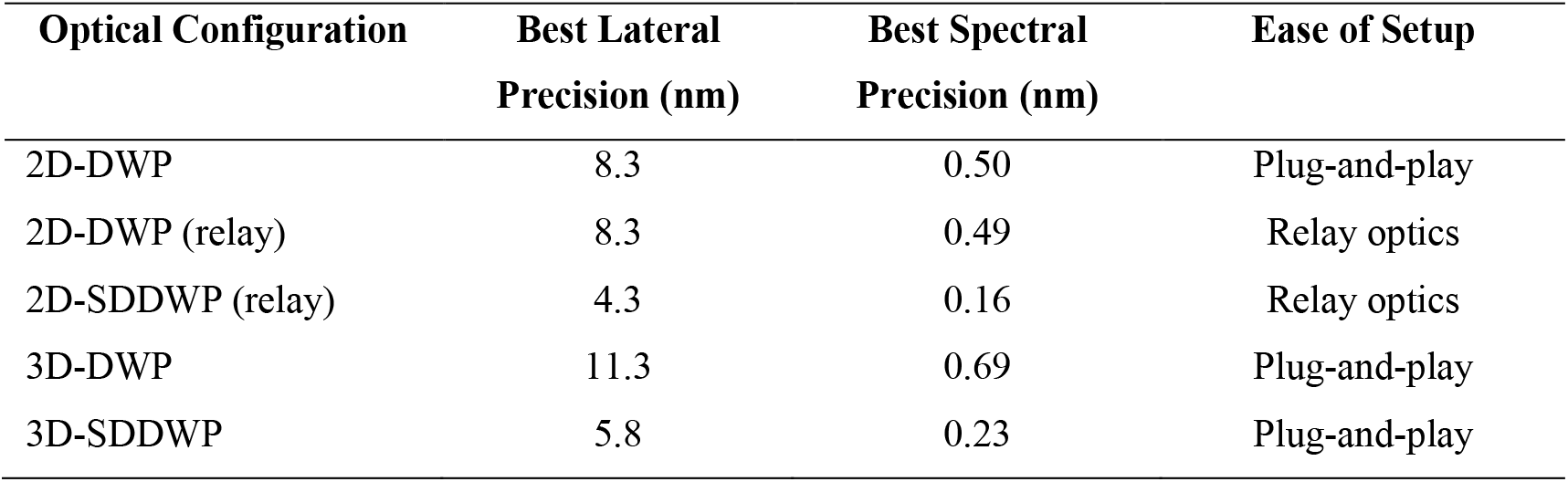
Summary of performance metrics for DWP–based sSMLM configurations.

## 4 Conclusion

In this work, we systematically compared DWP-based sSMLM configurations for 2D and 3D imaging, with emphasis on lateral precision, spectral precision, and practical ease of implementation. The quantitative comparison summarized in Table 1 highlights the trade-offs between optical performance and experimental complexity across the different configurations.

Overall, the SDDWP configurations provide the best performance. In particular, 2D-SDDWP with relay optics achieves the highest lateral (4.3 nm) and spectral (0.16 nm) precisions.

At the same time, 3D-SDDWP substantially outperforms 3D-DWP in both lateral and spectral precision despite the additional constraints imposed by biplane imaging (Table 1). These gains arise from symmetric dispersion, which increases effective spectral separation and improves photon utilization for localization by averaging information from paired diffraction orders.

However, 2D-DWP offers the simplest and most practical solution for imaging applications that do not require axial information. As shown in Table 1, 2D-DWP provides robust lateral (8.3 nm) and spectral (0.5 nm) precision in a compact, plug-and-play format. Its ease of setup and minimal alignment requirements make it particularly well-suited for routine 2D sSMLM experiments where reliability and experimental simplicity are prioritized over maximum achievable precision. In addition, we ignored the additional photon loss induced by the added relay optics and used only photon counts at the detector to perform the study.

In summary, while SDDWP configurations deliver the highest overall spatial and spectral performance, 2D-DWP represents the most accessible and user-friendly option for 2D spectroscopic SMLM. These results provide clear guidance for selecting an appropriate configuration based on whether ultimate performance or experimental simplicity is the primary consideration.

## Code, Data, and Materials Availability

The data presented in this article are publicly available in GitHub at https://github.com/FOIL-NU/DWP-Performance-Analysis.

## Acknowledgments

The authors sincerely acknowledge the generous support from NIH grants R01GM140478 and U54CA268084. Wei-Hong Yeo is supported by the Christina Enroth-Cugell Graduate Research Award at Northwestern University.

Menglin Shi is a Ph.D. Candidate in Biomedical Engineering at Northwestern University, specializing in optics and simulation for super-resolution microscopy. Working in the Functional Optical Imaging Laboratory, she has shown exceptional proficiency in her research. Menglin is passionate about advancing our understanding of biological systems at the microscopic level and is committed to making a significant impact in her field.

Wei-Hong Yeo is a Ph.D. graduate in Biomedical Engineering from Northwestern University, specializing in optical design and computational modeling for advanced biomedical imaging and super-resolution microscopy. Working in the Functional Optical Imaging Laboratory, he develops integrated optical systems and analysis methods that bridge hardware and data processing, with the goal of translating novel imaging technologies into practical tools for biological research.

Cheng Sun is a Professor of Mechanical Engineering at Northwestern University. He received his B.S. and M.S. degrees from Nanjing University, and his Ph.D. degree from Penn State University. His research interests include photonics and phononic metamaterials, plasmonics, nano-imaging and nano-manufacturing.

Hao F. Zhang is a Professor of Biomedical Engineering at Northwestern University. He received his B.S. and M.S. degrees from Shanghai Jiao Tong University and his Ph.D. degree from Texas A&M University. His research interests include optical coherence tomography, super-resolution imaging, vision science, and genomics.

## References

1. E. Betzig et al., “Imaging Intracellular Fluorescent Proteins at Nanometer Resolution,” Science 313(5793), 1642–1645 (2006) [doi: 10.1126/science.1127344].

2. M. J. Rust, M. Bates, and X. Zhuang, “Sub-diffraction-limit imaging by stochastic optical reconstruction microscopy (STORM),” Nat Methods 3(10), 793–796 (2006) [doi: 10.1038/nmeth929].

3. B. Dong et al., “Super-resolution spectroscopic microscopy via photon localization,” Nat Commun 7(1), 12290 (2016) [doi: 10.1038/ncomms12290].

4. Z. Zhang et al., “Machine-learning based spectral classification for spectroscopic single-molecule localization microscopy,” Opt. Lett. 44(23), 5864 (2019) [doi: 10.1364/OL.44.005864].

5. K.-H. Song et al., “Three-dimensional biplane spectroscopic single-molecule localization microscopy,” Optica 6(6), 709 (2019) [doi: 10.1364/OPTICA.6.000709].

6. K.-H. Song et al., “Symmetrically dispersed spectroscopic single-molecule localization microscopy,” Light Sci Appl 9(1), 92 (2020) [doi: 10.1038/s41377-020-0333-9].

7. Y. Suzuki et al., “Imaging of the fluorescence spectrum of a single fluorescent molecule by prism-based spectroscopy,” FEBS Letters, 5 (2002).

8. Z. Zhang et al., “Ultrahigh-throughput single-molecule spectroscopy and spectrally resolved super-resolution microscopy,” Nat Methods 12(10), 935–938 (2015) [doi: 10.1038/nmeth.3528].

9. M. J. Mlodzianoski et al., “Super-Resolution Imaging of Molecular Emission Spectra and Single Molecule Spectral Fluctuations,” PLoS ONE 11(3), M. Sauer, Ed., e0147506 (2016) [doi: 10.1371/journal.pone.0147506].

10. K.-H. Song et al., “Monolithic dual-wedge prism-based spectroscopic single-molecule localization microscopy,” Nanophotonics 11(8), 1527–1535, De Gruyter (2022) [doi: 10.1515/nanoph-2021-0541].

11. W.-H. Yeo, C. Sun, and H. F. Zhang, “Physically informed Monte Carlo simulation of dual-wedge prism-based spectroscopic single-molecule localization microscopy,” JBO 29(S1), S11502, SPIE (2023) [doi: 10.1117/1.JBO.29.S1.S11502].

12. B. Brenner et al., “Implementation and calibration of spectroscopic single-molecule localization microscopy,” BMC Methods 2(1), 2 (2025) [doi: 10.1186/s44330-025-00022-x].

13. W.-H. Yeo et al., “Maximizing photon utilization in spectroscopic single-molecule localization microscopy using symmetrically dispersed dual-wedge prisms,” npj Imaging 4(1), 20, Nature Publishing Group (2026) [doi: 10.1038/s44303-026-00152-z].

14. B. Rieger and S. Stallinga, “The Lateral and Axial Localization Uncertainty in Super-Resolution Light Microscopy,” ChemPhysChem 15(4), 664–670 (2014) [doi: 10.1002/cphc.201300711].

15. J. A. Kurvits, M. Jiang, and R. Zia, “Comparative analysis of imaging configurations and objectives for Fourier microscopy,” J Opt Soc Am A Opt Image Sci Vis 32(11), 2082–2092 (2015) [doi: 10.1364/JOSAA.32.002082].

